# A zero-inflated gamma model for post-deconvolved calcium imaging traces

**DOI:** 10.1101/637652

**Authors:** Xue-Xin Wei, Ding Zhou, Andres Grosmark, Zaki Ajabi, Fraser Sparks, Pengcheng Zhou, Mark Brandon, Attila Losonczy, Liam Paninski

**Affiliations:** Departments of Statistics and Neuroscience, Grossman Center for the Statistics of Mind, Center for Theoretical Neuroscience, and Zuckerman Mind Brain Behavior Institute, Columbia University; Department of Neuroscience, Zuckerman Mind Brain Behavior Institute, Columbia University; Integrated Program in Neuroscience, McGill University; Douglas Hospital Research Centre, McGill University

## Abstract

Calcium imaging is a critical tool for measuring the activity of large neural populations. Much effort has been devoted to developing “pre-processing” tools applied to calcium video data, addressing the important issues of *e.g.*, motion correction, denoising, compression, demixing, and deconvolution. However, computational modeling of deconvolved calcium signals (i.e., the estimated activity extracted by a pre-processing pipeline) is just as critical for interpreting calcium measurements. Surprisingly, these issues have to date received significantly less attention. To fill this gap, we examine the statistical properties of the deconvolved activity estimates, and propose several density models for these random signals. These models include a zero-inflated gamma (ZIG) model, which characterizes the calcium responses as a mixture of a gamma distribution and a point mass which serves to model zero responses. We apply the resulting models to neural encoding and decoding problems. We find that the ZIG model out-performs simpler models (e.g., Poisson or Bernoulli models) in the context of both simulated and real neural data, and can therefore play a useful role in bridging calcium imaging analysis methods with tools for analyzing activity in large neural populations.

## 1 Introduction

Calcium imaging is one of our primary methods for measuring activity of large neural populations at single-cellular resolution (Yuste and Denk, 1995; Svoboda et al., 1997; Helmchen et al., 1999; Dombeck et al., 2010). Calcium imaging has several well-known advantages: this approach offers sub-cellular spatial resolution with cell-type specificity and can be coupled easily with a variety of genetic tools; the method has proven scalability to record simultaneously from thousands of neurons in vivo; finally, it allows for longitudinal tracking of cellular activity across multiple days.

At the same time, calcium imaging presents some important analysis challenges: calcium signals represent a slow, nonlinear encoding of the underlying spike train signals of interest, and therefore it is necessary to denoise and temporally deconvolve temporal traces extracted from calcium video data into estimates of neural activity. These issues have received extensive attention in the literature (Vogelstein et al., 2009; Vogelstein et al., 2010; Pnevmatikakis et al., 2016; Deneux et al., 2016; Theis et al., 2016; Friedrich et al., 2017; Jewell et al., 2018; Speiser et al., 2017; Aitchison et al., 2017; Berens et al., 2018; Pachitariu et al., 2018; Greenberg et al., 2018). Some of these deconvolution approaches estimate spiking probabilities directly (Vogelstein et al., 2009; Pnevmatikakis et al., 2016; Deneux et al., 2016; Speiser et al., 2017; Aitchison et al., 2017; Greenberg et al., 2018), but many approaches instead estimate the influx of calcium in each time bin, rather than a spiking probability (Vogelstein et al., 2010; Pnevmatikakis et al., 2016; Friedrich et al., 2017; Jewell et al., 2018; Berens et al., 2018; Pachitariu et al., 2018; Stringer and Pachitariu, 2019); these non-probabilistic approaches tend to be faster and are therefore popular in practice.

What is a proper statistical model for the output of these non-probabilistic calcium deconvolution approaches? Somewhat surprisingly, this question has received relatively less attention. The simplest approach is to simply threshold the deconvolved output and treat the resulting super-threshold events as “spikes,” but this approach clearly discards information about the number of spikes per bin, and there is no obvious optimal way to set the threshold. Another naive approach would be to apply standard point-process models (e.g., Poisson regression models) to the deconvolved output — but as we will see below, the Poisson model is a poor approximation here, not least because the deconvolved output can take continuous values, while the Poisson distribution is supported on the integers.

In this paper, we investigate statistical models to characterize the deconvolved calcium activity. In particular, we propose a zero-inflated gamma (ZIG) model, a two-component mixture model including a “spike” of probability at zero response and another continuous component for modeling positive responses, specified by a gamma distribution. We apply this model to simulated data and real imaging datasets from hippocampus and thalamus, and find that it provides good fits across a wide variety of deconvolution parameters and data types. Next we show that the ZIG model can be embedded within “encoding models” to characterize the probability of calcium responses given time-varying covariates such as the location or orientation of the animal during behavior. Finally, we demonstrate that the ZIG-based encoding model leads to more accurate Bayesian decoding of these covariates.

## 2 Results

### 2.1 Nonnegative deconvolution methods applied to calcium fluorescence traces produce a mixture of zeros and positive real-valued output, well-captured by the zero-inflated gamma model

We begin with simple simulated data (Figure 1a-d): we generate a Poisson spike train, then push this spike train through a standard auto-regressive AR(1) model for calcium response (Vogelstein et al., 2010) and add noise to generate simulated fluorescence traces, and then run a popular non-negative deconvolution method (Vogelstein et al., 2009; Friedrich et al., 2017) to obtain the post-deconvolution response, denoted as 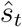. With experimentally relevant signal-to-noise levels, the resulting histogram of deconvolved responses 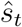 typically has a “spike-and-slab” form (Figure 1b-d, right): significant mass is placed exactly at zero (the “spike”), with the remaining mass forming a continuous “slab” on the positive real axis. (This spike-and-slab structure of 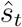 is unsurprising: the deconvolution approach applied here enforces sparsity and non-negativity constraints on 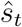, forcing 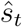 to be exactly zero for many timesteps *t*.) Empirically, a shifted gamma model^1^ suffices to capture the shape of the slab (green traces in Figure 1b-d, right); the shift is fixed to be equal to the minimum spike size allowed by the deconvolution algorithm (“*s*_*min*_”), and therefore the gamma distribution is still specified by two parameters. We denote the resulting three-parameter distribution (with the third parameter corresponding to the probability of a non-zero response) as the “zero-inflated gamma” (ZIG) model.

**Figure 1:**
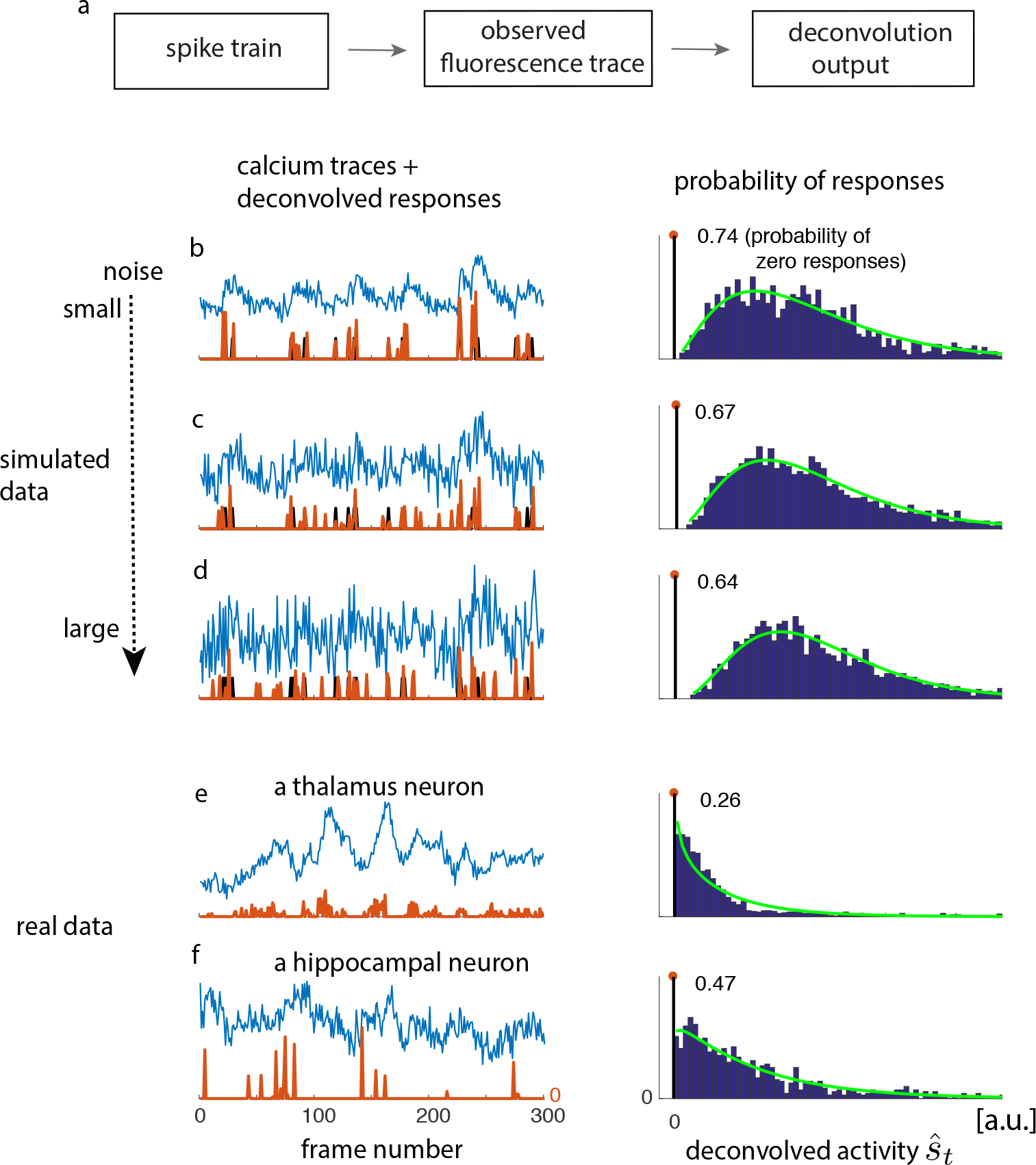
Illustration of the zero-inflated gamma (ZIG) model: deconvolved calcium responses typically consist of a mixture of zero responses plus a continuous component that is well-modeled as a gamma distribution, in both simulated and real imaging data. (a) Pipeline of the simulations for the generation of artificial data. The spike train is sampled from a homogeneous Poisson process. The calcium concentration is then determined by a auto-regressive process (AR(1) process, with decay time constant 450ms), driven by the spike train. The observed calcium trace is determined by the calcium concentration plus independent Gaussian noise. The deconvolved calcium responses are obtained using the OASIS deconvolution algorithm described in (Friedrich et al., 2017). (b,c,d) left: observed fluorescence trace (blue), ground truth spikes (black), and the deconvolved output (orange). Each frame = 30ms. Right: the histogram of the deconvolved output (blue) and the ZIG fit (green); the number on each histogram represents the proportion of zero responses. The additive noise level of the simulated fluorescence increases from panel b to panel d. (e) Observed fluorescence and deconvolved response of a neuron from ADN (see appendix for full experimental details). (f) Same as panel e but from the hippocampus (again, see appendix for full experimental details). (Conventions as in b-d but we no longer have access to the true spikes.) In each case, the ZIG model provides a good fit to the deconvolved outputs.

In these simulations, we have made several simplifying assumptions, including: i) a simple AR(1) model for the generative process of the calcium fluorescence; ii) the a priori knowledge about the time constant of the AR process; iii) the increase of calcium concentration following each spike has a constant, deterministic size. Presumably, any deviations from these assumptions would make the estimated “spikes” 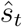 noisier, thus making the continuous part of the response histogram smoother. On the other hand, increasing the signal-to-noise ratio (SNR) and/or making the neurons burstier can introduce multiple “bumps” in the continuous part of the distribution (not shown). This multiple-bump case could potentially be handled by incorporating a multiple-component mixture model for the slab in our spike-and-slab model, but (as we will discuss next), in practice for real data we have not found this to be necessary and have not pursued this direction systematically.

We turn next to real data. In most real datasets, the ground truth spiking (the first part of our simulation pipeline outlined in Figure 1a) is not available, but nonetheless we can run the same deconvolution algorithm on the observed fluorescence trace to obtain 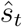. The resulting histogram of 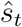 is again well-fit by the ZIG model, for two example neuron cases shown in Figure 1e-f.

### 2.2 The ZIG model is applicable to the outputs of multiple deconvolution methods, applied to data from multiple calcium indicators

We next seek to determine whether the observations made in Figure 1 are specific to a particular deconvolution method or calcium indicator.

In Figure 2, we examine three deconvolution methods, including an *L*_1_-penalized method with a soft threshold (Vogelstein et al., 2010; Pnevmatikakis et al., 2016), a method with a hard threshold (i.e., positive minimal spike size “*s*_*min*_”) (Friedrich et al., 2017), and an *L*_0_-penalized method (Jewell and Witten, 2018; Jewell et al., 2018). Each method has a free parameter which controls the sparsity of the inferred responses; varying this parameter leads to corresponding changes in the histograms of the deconvolved responses 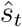, with more or less probability mass assigned to 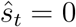. Over a range of parameters, the ZIG model provides a good fit to the output histogram for all three of the algorithms examined here.

**Figure 2:**
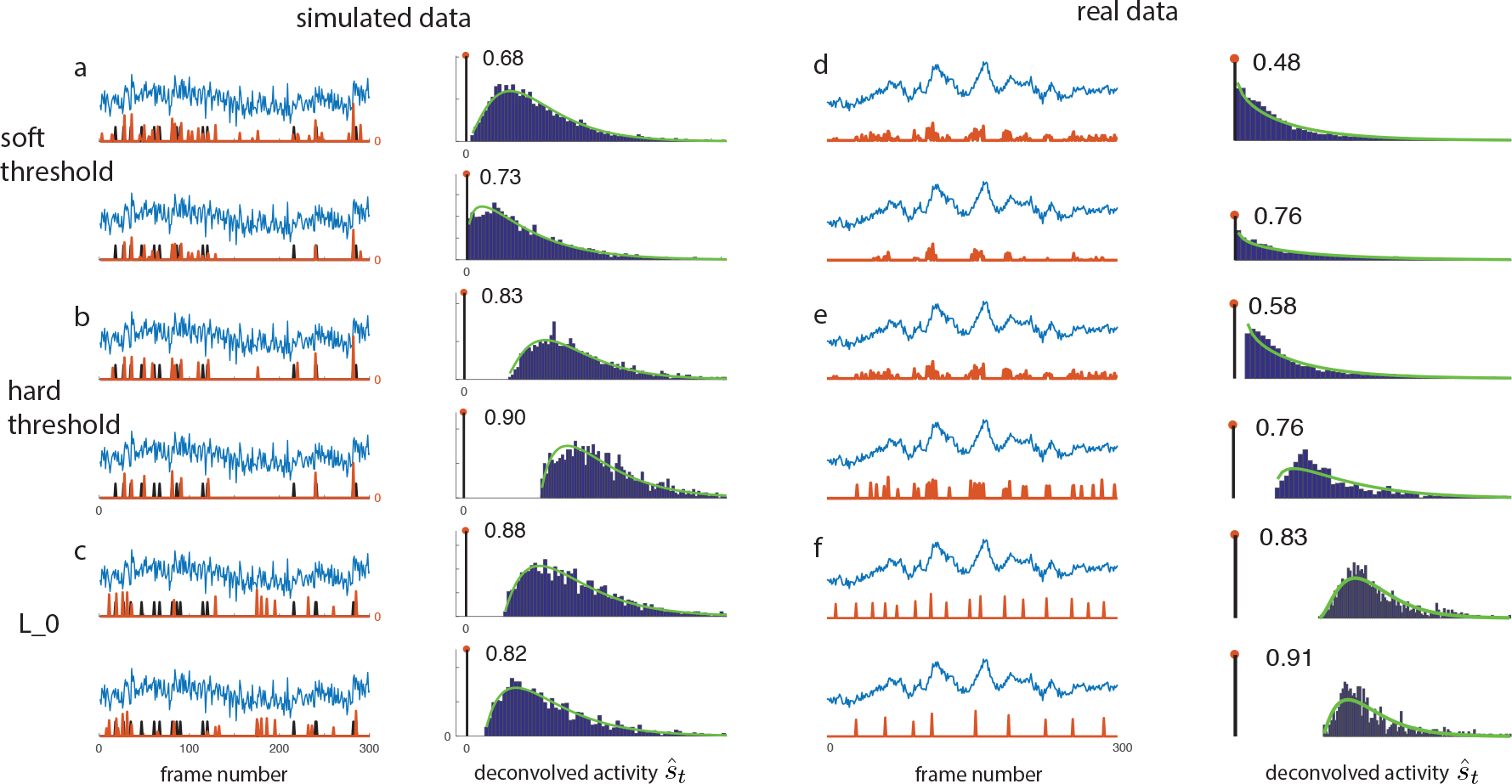
The ZIG model is robust with respect to the details of different deconvolution methods. We consider three deconvolution methods, including a *L*_1_-penalization method with soft threshold (panel a,d) (Pnevmatikakis et al., 2016), a method with non-zero minimal spike size or hard threshold (panel b,e) (Friedrich et al., 2017), and an *L*_0_-penalized method (panel c,f) (Jewell et al., 2018). We apply these methods to both simulated (panel a,b,c) and real data (panel d,e,f; same traces as in Figure 1). Each method has a hyper-parameter controlling the sparseness of the deconvolved activity. Two values of the sparseness parameter are examined for each method. Conventions as in Figure 1.

Next, in Figure 3, we examine data shared through the SpikeFinder challenge (Berens et al., 2018), including traces recorded using four calcium indicators (GCamp6s, jRCAMP1a, OGB-1, jRGECO1a). Again, we find that the ZIG model provided a good fit across a wide range of data.

**Figure 3:**
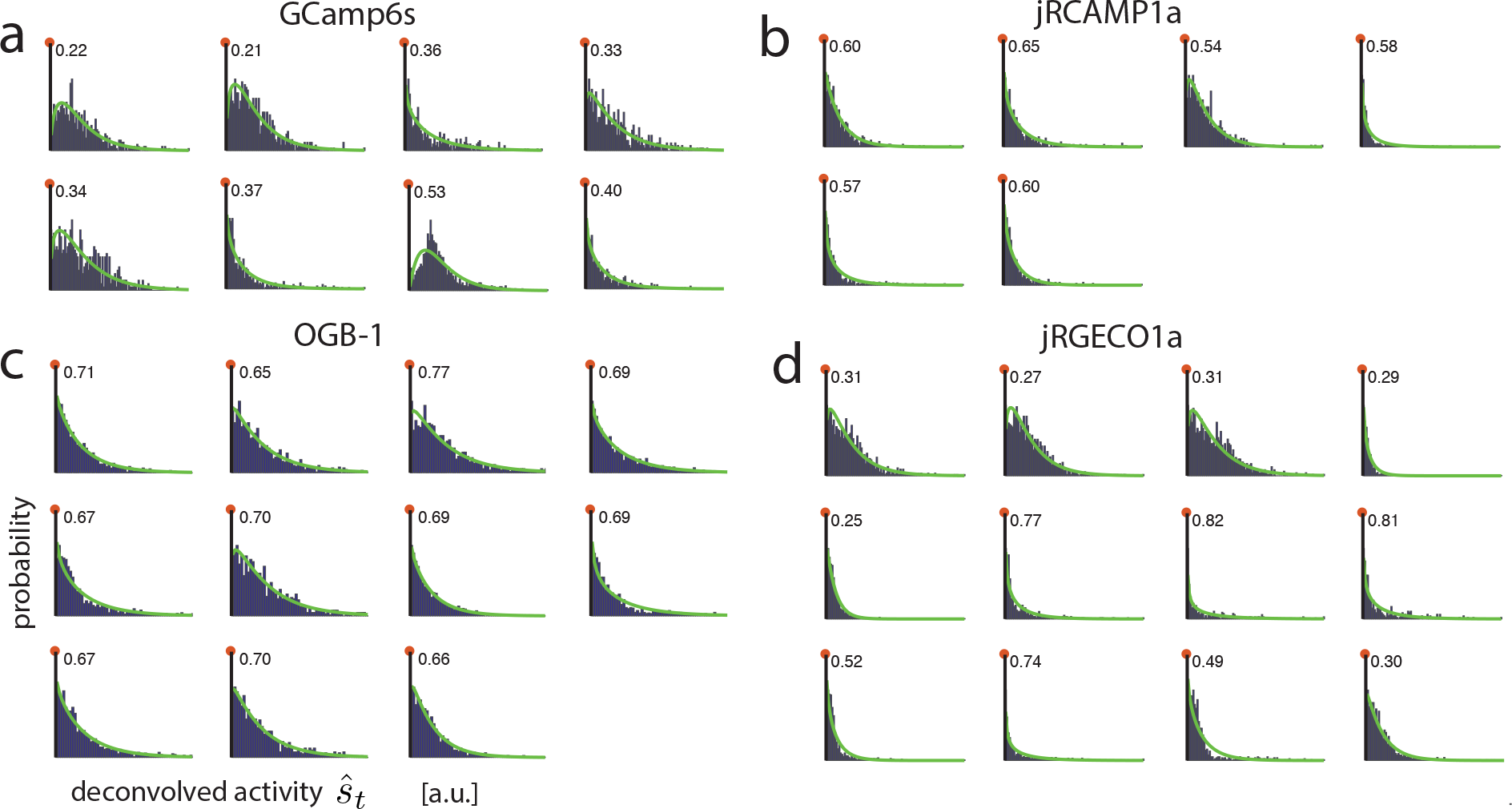
The ZIG model is robust with respect to data collected with different calcium indicators. Data collected using four different calcium indicators are tested. The data are from the SpikeFinder challenge dataset (panel a,c from (Theis et al., 2016); panel b,d from (Dana et al., 2016)). (Specifically, the four datasets used here are SpikeFinder dataset #1 (calcium indicator OGB-1), dataset #3 and #5 (indicator GCamp6s), dataset #9 (indicator jRCAMP1a), and dataset #10 (indicator jRGECO1a).) All the neurons examined here fired at least 200 spikes.

### 2.3 Constructing encoding models for simulated calcium responses

We have seen above that the ZIG model provides a good fit to the marginal distribution of the deconvolved responses 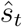. Now we want to exploit this probabilistic model to perform neural data analysis tasks. The first step is to fit *encoding models*: ie, what is 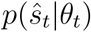, for some observed covariate *θ*_*t*_ such as a stimulus or movement. In general, *θ*_*t*_ may be multi-dimensional, but in the example applications here *θ*_*t*_ will be one dimensional. Once these encoding models are fit and validated, we can use them to perform tasks like decoding of *θ*_*t*_ given the observed deconvolved responses 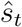. The overall approach is illustrated in Figure 4.

**Figure 4:**
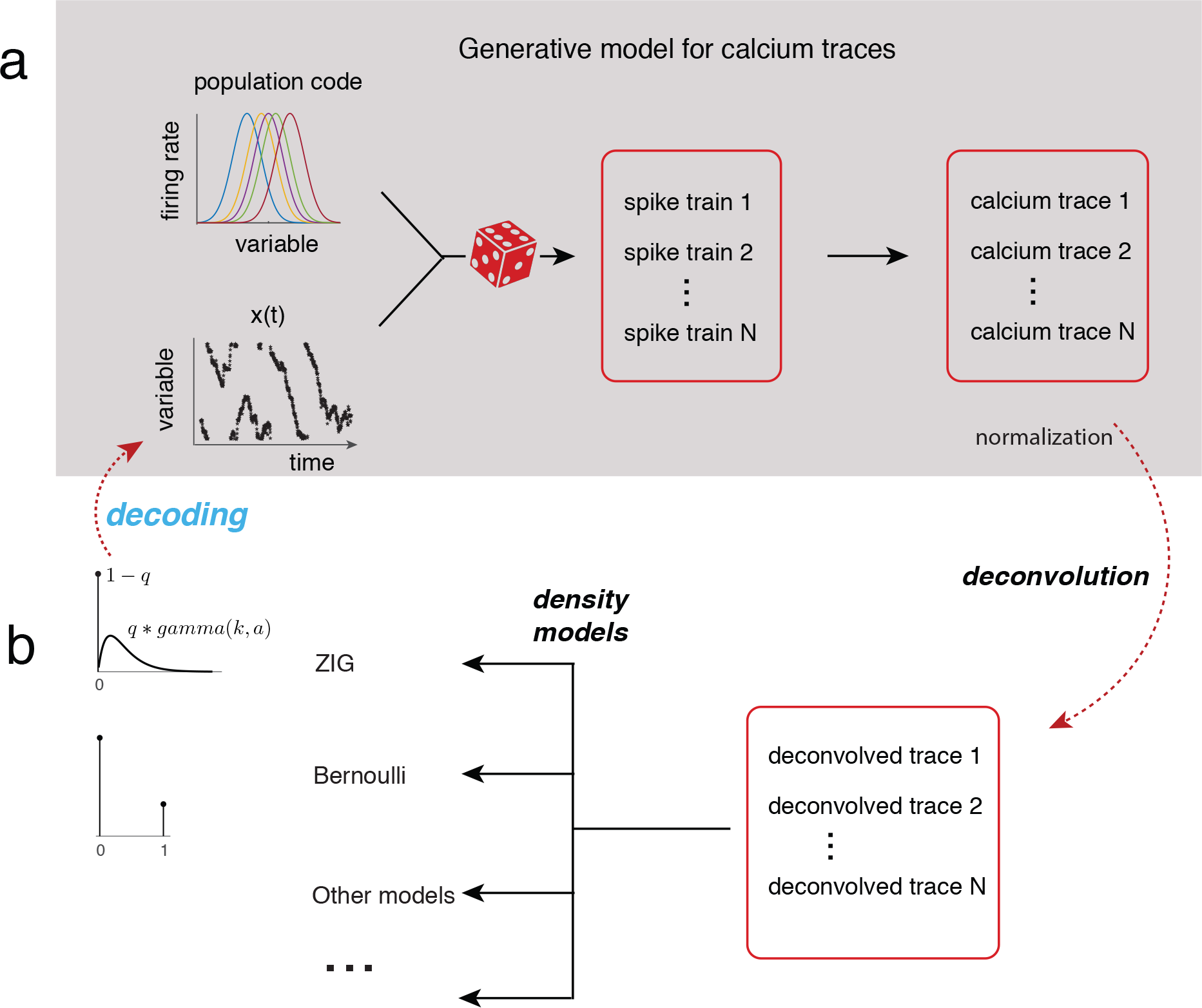
The encoding-decoding modeling framework. (a) The full generative “encoding” model for calcium fluorescence traces. Given the neural tuning curves and the covariate sequence *θ*_*t*_, spikes are generated probabilistically and transformed (as in Figure 1) into the observed fluorescence traces. (b) After deconvolving to obtain 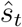, we can use the ZIG as well other models for 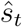, and use the estimated encoding model and 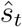 to decode *θ*_*t*_.

To fit the ZIG model to 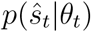, we need to fit three parameters, each of which may depend on *θ*: the probability of non-zero response *q*(*θ*), the scale parameter *a*(*θ*), and the shape parameter *k*(*θ*) for the gamma component, specifically

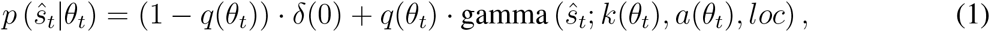

where we fix the location parameter *loc* for the gamma component as the minimum spike size “smin” (The mean of the ZIG model with parameters (*q*, *k*, *a*, *loc*) is *q*(*ka* + *loc*), and the variance is *qka*^2^ + *q*(*ka* + *loc*)^2^(1 − *q*).) We model these parameters as nonlinear functions of *θ*_*t*_; we use neural networks to parameterize these nonlinearities, and then estimate the weights of these networks by maximum likelihood (see appendix for full details).

To test this approach, we generate artificial calcium imaging datasets with hundreds of simulated neurons (Figure 4). We first construct tuning curves of individual neurons that tile the space of *θ* values. In the real data examples presented below, *θ* will be a one-dimensional variable (e.g., the animal’s head direction), so we use a one-dimensional *θ* in these simulations. Next we take a empirically measured time series *θ*_*t*_(the head direction of a mouse), and compute the time-varying firing rates for individual neurons by plugging *θ*_*t*_ into the tuning curves. We then generate binned spike trains with different noise characteristics; we experiment with spike counts drawn from a Poisson distribution or a negative binomial (NB) distribution, as both have been proposed to model empirically observed spike responses (Tomko and Crapper, 1974; Tolhurst et al., 1981; Goris et al., 2014). Next we plug these simulated binned spike trains into the same generative model for calcium fluorescence traces discussed above, then deconvolve the resulting traces to obtain simulated responses 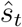. Finally, we fit the ZIG encoding model to the resulting responses 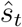.

We compare the ZIG model against simpler Poisson, Bernoulli, and gamma models (see appendix for full details). Figure 5 shows the results from a simulated dataset with negative binomial spiking. (We find that the results on the Poisson dataset are qualitatively similar; data not shown.) Overall, for both the Poisson or NB simulated datasets, we find that all of these models except for the Bernoulli model can capture the data mean well (the Bernoulli model is only effective for data in which the mean of 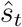 in each bin is bounded below 1; this model fails to capture the responses in bins with high firing rates). However, only the ZIG model can properly capture both the mean and variability of 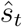. The alternative models tend to either over- or under-estimate the variance, therefore providing poor descriptions of the distributions of the deconvolved responses; thus, the extra flexibility (due to the larger number of parameters) in the ZIG model is necessary to capture basic statistics of the data.

**Figure 5:**
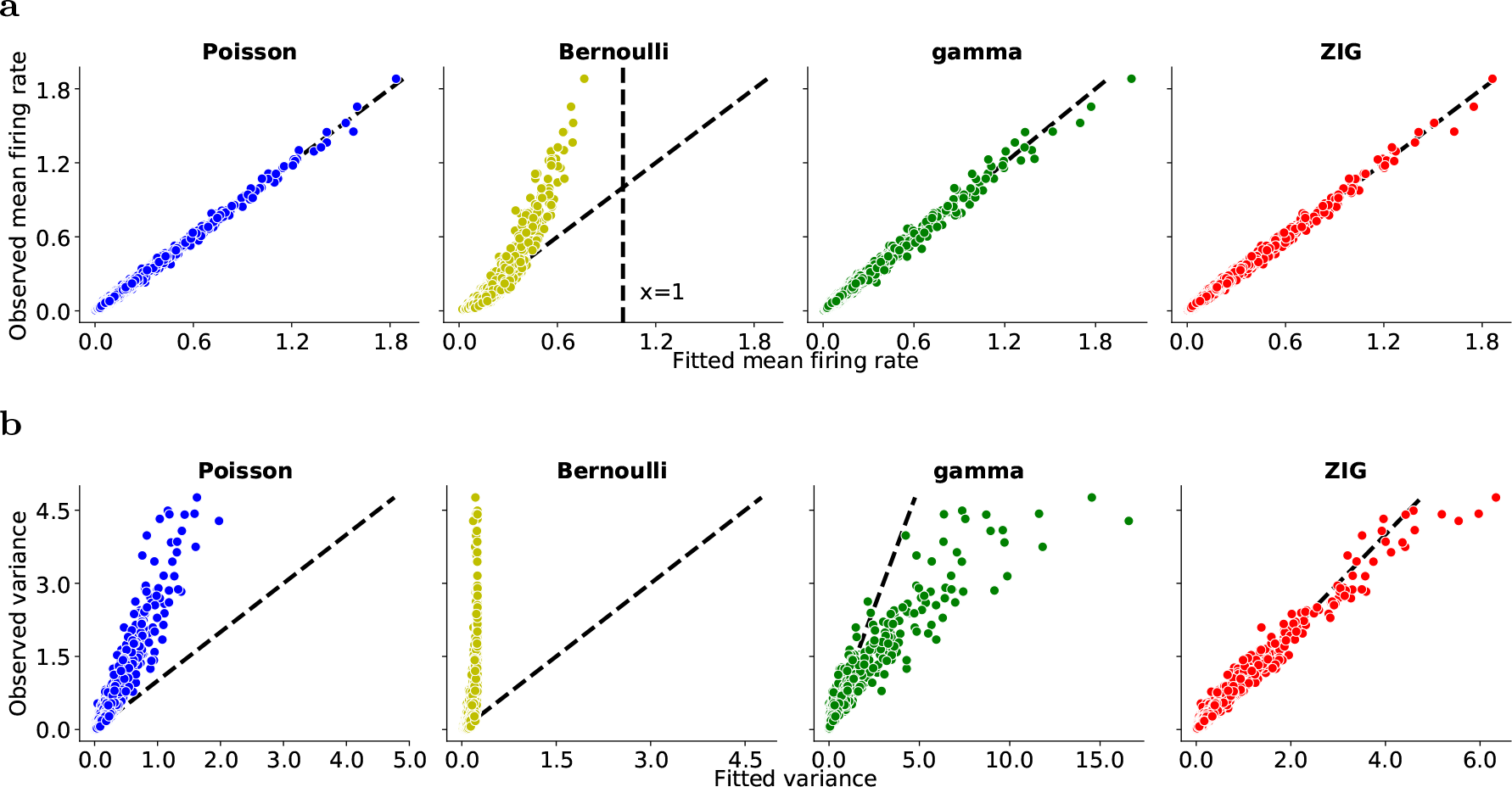
The ZIG model captures the means and variances of deconvolved calcium responses in the simulated data. (a,b) Summary plots based on all the neurons (*N* = 215) showing the observed versus the predicted mean/variance of 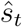. We divide the *θ* range to into small equi-spaced bins (number of bins = 18 here), and compute the mean and variance for the observed and fitted responses corresponding to each bin for each model neuron. Each dot represents the mean (or variance) associated with one bin from one neuron.

### 2.4 The ZIG encoding model leads to improved Bayesian decoding in simulated data

In the previous section we showed that the ZIG encoding model is flexible enough to capture the mean and variance of 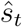 across a wide range of firing rate regimes, in simulated data. Can we exploit this encoding model to obtain an improved decoder for *θ*_*t*_? We use a classic Bayesian decoding approach to address this question: we compute the posterior distribution of *θ*_*t*_, under the different encoding models for 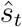 discussed above, and then quantify how well the resulting posterior distributions capture the uncertainty in *θ*_*t*_ given the observed 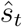.

We quantify the performance of the resulting decoders on simulated data in Figure 6. Overall, the ZIG model leads to the smallest decoding error over a wide range of deconvolution sparsity parameters. Interestingly, accuracy degrades monotonically as a function of the sparsity of the output 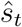: i.e., the decoders can take advantage of even very small outputs 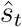 to improve the decoding accuracy. (In Figure 6 we use the deconvolution approach from (Friedrich et al., 2017), with a hard-threshold on the minimal spike size; results based on the soft-threshold deconvolution approach from (Pnevmatikakis et al., 2016) are similar.) The decoder based on the ZIG encoding model also achieves the highest coverage rate (i.e., the posterior credible interval covers the true value of *θ*_*t*_ with highest probability). In contrast, the decoders based on the Poisson and gamma models output credible intervals with mis-calibrated coverage rates (i.e., the credible interval based on these models was narrower than it should have been), due to a mismatch between the true versus the modeled distribution of 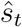. In other words, a Bayesian statistician using a Poisson or gamma encoding model would be (mistakenly) overly confident in her predictions.

**Figure 6:**
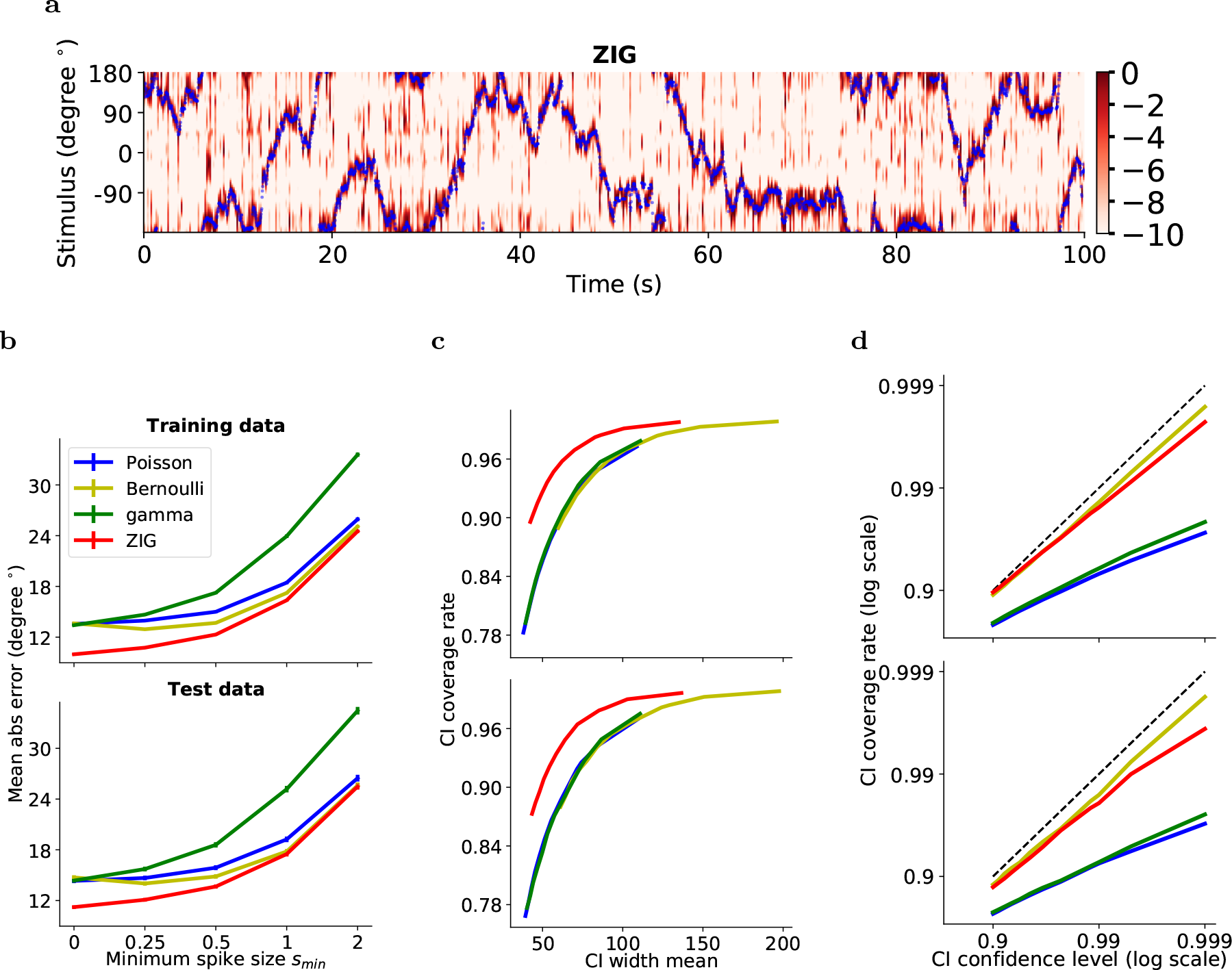
The ZIG model leads to improved decoding performance on simulated data. Decoding is performed based on the deconvolved responses in single frame, time window ~ 33ms, with no smoothing across frames on all neurons (*N* = 215). (a) True simulated location (blue) plotted with the decoded normalized log-posterior probability under the ZIG model (red; posterior at each step is normalized to have a maximum of 1, for easier visualization). Note that the decoded posterior does a good job of tracking the true location. (b) Decoding mean absolute error ±1 standard error under different encoding models, with varying *s*_*min*_, the minimum spike size parameter in (Friedrich et al., 2017); larger values of *s*_*min*_ correspond to sparser output 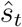. (c) Posterior credible interval (CI) width vs CI coverage rate (the probability that the true location falls within the CI; higher is better here). (d) Confidence level vs CI coverage rate. Dashed line indicates unity (i.e., the CI is achieving its nominal coverage rate). In (b,c) we see that the ZIG model leads to the lowest decoding error and the best coverage rate over a range of parameters, while (d) shows that the CI computed under the ZIG model achieves a nearly-nominal coverage rate, as desired; in contrast, the Poisson and gamma encoding models output mis-calibrated credible intervals.

### 2.5 Application to real imaging data

In the previous section we developed the encoding-decoding analysis pipeline on simulated data. Next we apply these methods to real data. We focus on two calcium imaging datasets in this section. The first is a single-photon imaging dataset collected from thalamic region ADN, and the second is a two-photon dataset from hippocampal region CA1. Both datasets are collected in animals performing spatial navigation tasks (see Methods for full details). Our aim is to decode head direction (during free behavior) in the ADN data and location along a circular track (during head-fixed behavior) in the hippocampal data; thus in both cases the variable *θ* is one-dimensional, as in the simulated data.

We begin in Figure 7 by fitting encoding models to the ADN data. The results are similar to those shown in Figure 5: of the models examined here, only the ZIG model can capture both the mean and the variance of the empirical data. Further, in panel c we examine the tuning curves from this population of neurons. We compute the mean firing rates as a function of *θ* (leftmost panel) and plot these next to the estimated *a*(*θ*) and *q*(*θ*) curves (middle and right panels); recall that the mean of 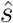 as a function of *θ* scales proportionally with *a*(*θ*)*q*(*θ*) in the ZIG model. We see that the parameters *a*(*θ*) and *q*(*θ*) covary across this population, indicating that there may be some statistical benefit in fitting these parameters with a hierarchical model that can share information between *a*(*θ*) and *q*(*θ*); however, we have not pursued this direction systematically.

**Figure 7:**
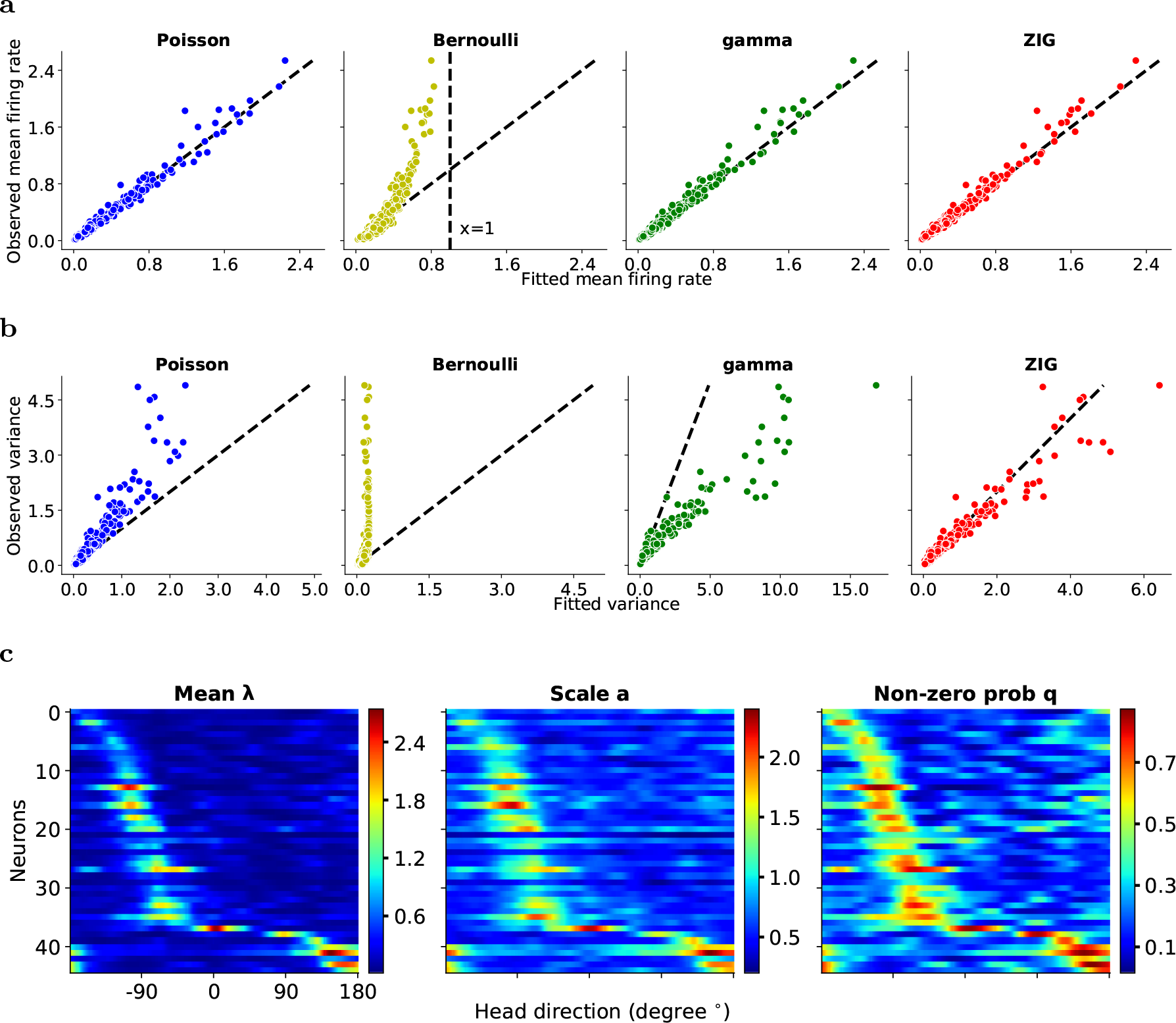
The ZIG model best captures the means and variances of deconvolved calcium responses in the ADN dataset. (a,b) Conventions as in Figure 5; similar to results based on the simulated data, the ZIG model again provides the best fits to the observed means and variances for this experimental dataset. (c) Estimated parameter values for the ZIG model and Poisson model, for multiple neurons (each row corresponds to one neuron, while the columns correspond to different *θ* values). Neurons are sorted according to the preferred firing direction. For the Poisson model, each row plots the mean firing rate λ as a function of head direction for each neuron. For the ZIG model, there are three sets of parameters. The scale parameter *a* and the probability of non-zero response *q* are plotted. Notice that the two parameters are correlated; both parameters tend to scale with the estimated mean rate λ. We find that we could obtain good fits by fixing the shape parameter *k* in the ZIG model as a function of *θ* (data not shown).

**Figure 8:**
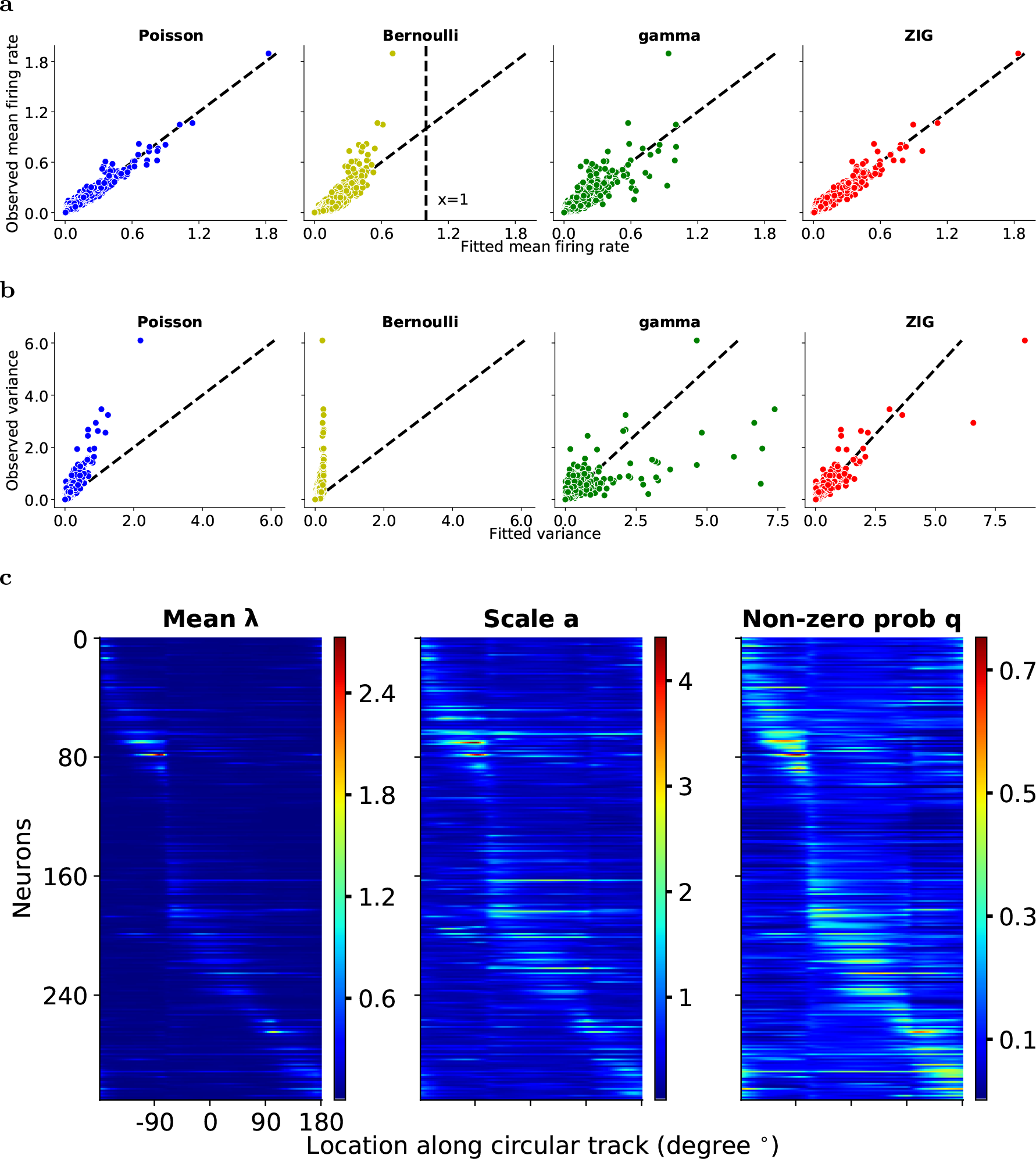
Encoding results for hippocampus data during running. Conventions as in Figure 7. Again, among the four models considered, only the ZIG model can capture both the mean and variance of the calcium responses conditional on the animal’s location on the track.

Next we turn to decoding (Figures 9 and 10). Again, the results of the real data analysis are largely consistent with the simulated results presented in Figure 6: in both datasets, the ZIG encoding model leads to more accurate Bayesian decoding, with higher credible interval coverage rates. Again, the decoding accuracy improves as the sparsity of 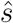 decreases.

**Figure 9:**
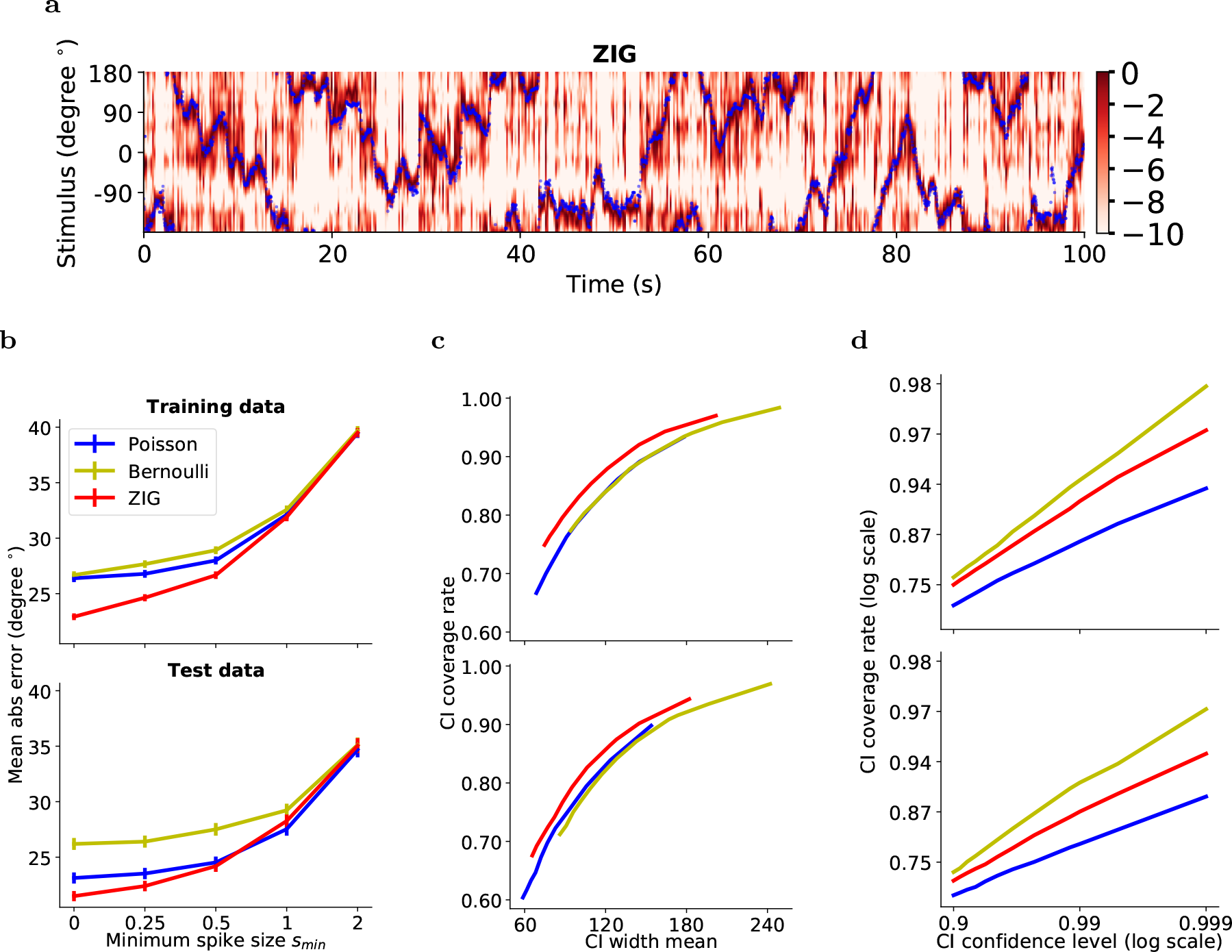
Decoding results for head direction (ADN) dataset. Decoding is performed based on responses from single frame, time window ~ 33ms. Conventions similar to Figure 6. The gamma model performed poorly here and is not shown.

**Figure 10:**
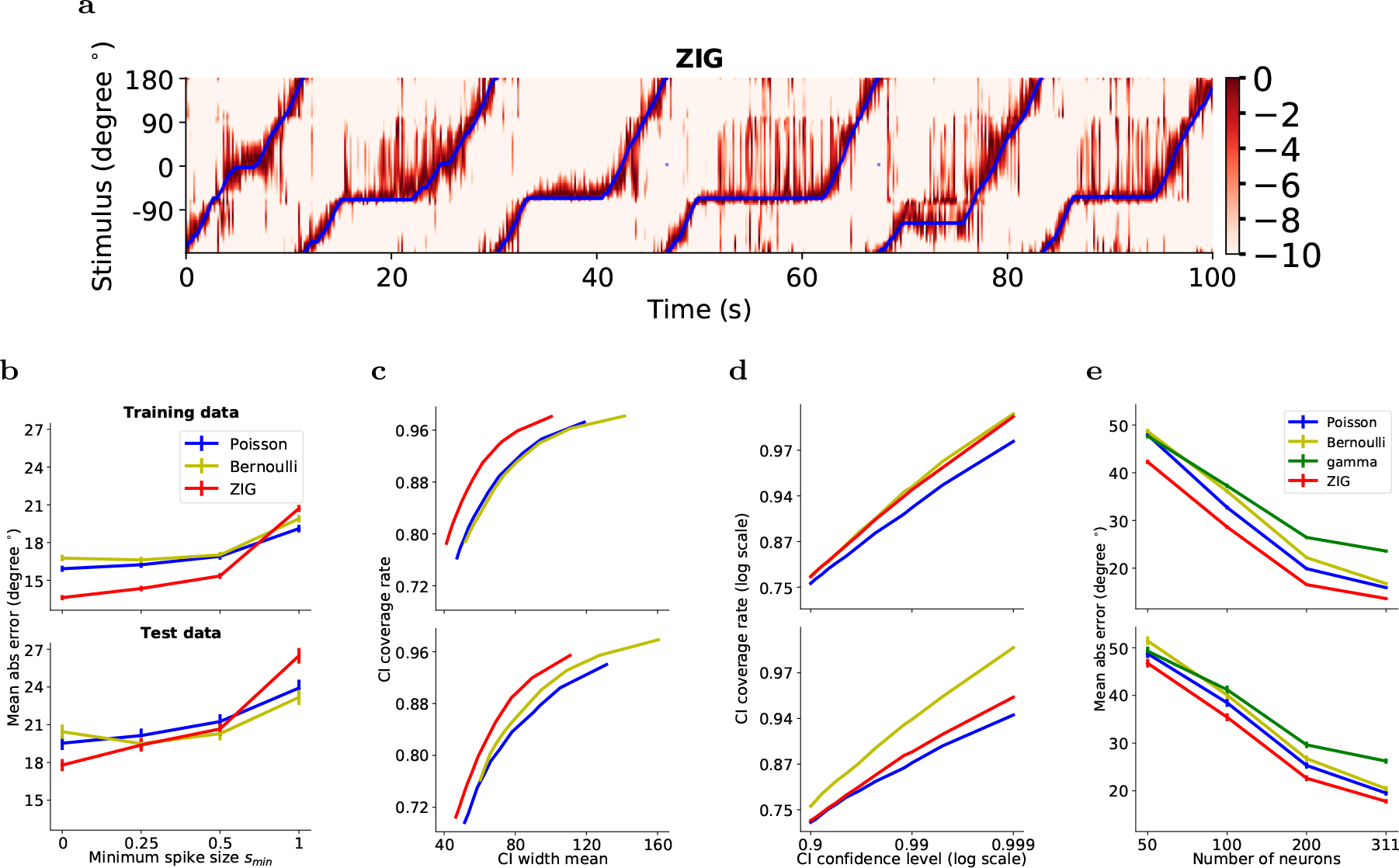
Decoding results for hippocampus data. The encoding model is estimated based on the data during running, as is typical in the hippocampal decoding literature (Davidson et al., 2009). Decoding time window is set to be every two frames, ~ 33ms. (a,b,c,d) similar to Figure 9. (e) Decoding results for subsets of neurons of hippocampus data. We randomly select neurons (*N* = 50, 100, 200) out of the 311 total neurons and perform the encoding-decoding analysis based on these subsets. The advantage of the ZIG model remains robust with smaller sub-populations of hippocampal neurons.

One major difference between the simulated and real data is that the coverage probabilities are no longer well-calibrated, for any of the encoding models. In other words, the Bayesian posterior based on these encoding models is overly confident. We believe this is due to model mismatch: in our Bayesian decoder we model the responses 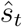 as conditionally independent across neurons given *θ*_*t*_, i.e.,

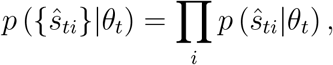

where 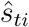 denotes the observed response at time *t* from cell *i*. This assumption makes a testable prediction: 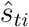 should be uncorrelated with 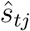 (where *i* and *j* index two different neurons) if we restrict attention to responses within a single bin of *θ* values. We find that these “noise correlations” are empirically not zero (invalidating the conditional independence assumption), and in fact if we perform a shuffling analysis in which we randomize the responses 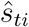 within each *θ* bin (thus preserving the relationship between 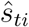 and *θ*_*t*_ while destroying noise correlations between 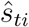 and 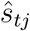), then we find that the calibration of the credible interval is restored (data not shown). We leave further detailed modeling of these noise correlations to future work.

## 3 Conclusion

The primary conclusion of this work is that the ZIG model provides a significantly improved fit to the distribution of the post-deconvolved calcium responses 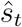: the ZIG model is sufficiently flexible to capture the zero-inflation and varying mean and dispersion of the data across a wide variety of indicators, deconvolution methods, and behavioral settings. Moreover, it is straightforward to extend this into a *θ*_*t*_-dependent encoding model, and in turn to use this encoding model for Bayesian decoding. The improved encoding fits provided by the ZIG lead directly to more accurate decoding, with better-calibrated posterior uncertainties. Finally, somewhat surprisingly, we find that setting the deconvolution hyperparameter to minimize the sparsity of 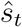 consistently leads to the most accurate decoder (consistent with results in (Pachitariu et al., 2018)); i.e., attempting to discard small “noisy spikes” in 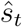 may be counterproductive.

Open source code implementing these methods is available at https://github.com/zhd96/zig. We hope these methods will be useful for the variety of downstream analyses that are currently being pursued by the calcium imaging community.

## Acknowledgments

We gratefully acknowledge support from NIBIB R01 EB22913 (LP), NSF Neuronex DBI-1707398 (LP), NIH 1U19NS104649-01 (LP), R01 NS094668 (AL), U19 NS104590 (AL), R01 MH100631 (AL), Charles H. Revson Foundation Biomedical Fellowship (AG), National Sciences and Engineering Council Discovery grant #74105 (MB), the Canada Research Chairs Program (MB), the Gatsby Charitable Foundation to the Center for Theoretical Neuroscience at Columbia University. We thank Tian Zheng and John Cunningham for helpful discussions.

## Supplementary Information

### 4 Materials and Methods

#### 4.1 Density models of the deconvolved calcium trace

##### ZIG model

We model the density of the devonvolution output as

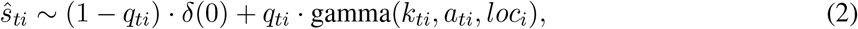

where *q*_*ti*_ denotes the probability of non-zeros, *a*_*ti*_ is the scale parameter of the gamma distribution, and *k*_*ti*_ is the shape parameter of the gamma distribution, for neuron *i* and time *t*. *loc*_*i*_ is the location parameter of the gamma distribution for neuron *i*, fixed as the minimum spike size *s*_*min*_. We denote *a*_*i*_ = (*a*_1*i*_, …,*a*_*Ti*_)^*T*^ ∈ 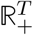 as the scale parameters for neuron *i*. Parameters *q* and *k* are defined similarly. We denote this family of density functions as 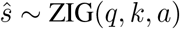. Note that when *s*_*min*_ = 0 the ZIG family has a useful scale-invariance property: if 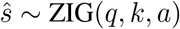, then 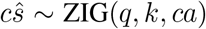, for any constant *c* > 0. This is convenient because in general the scale factor connecting spikes to increases in calcium concentration is unknown (and will typically vary from cell to cell); however, the scale invariance of the ZIG model implies that we never need to estimate this scale factor explicitly.

##### (Scaled-)Poisson model

The Poisson model places all of its probability mass on the non-negative integers, and is therefore inappropriate for modeling 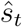, which has range 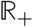. Nonetheless, as discussed below, it is possible to assign a pseudolikelihood under the Poisson model to real-valued observations 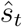, and to fit the Poisson rate λ by maximizing this pseudo-likelihood. However, the Poisson model does not have the scale-invariance property enjoyed by the ZIG model, and therefore some care must be taken in defining a scale for 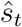 (empirically, we find that the performance of the Poisson encoding and decoding models are highly sensitive to scaling of 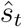). We experiment with two scaling approaches. In the first scheme, the deconvolved trace is normalized by the noise standard deviation of the raw calcium trace, using methods proposed previously (Pnevmatikakis *et al.*, 2016). In the second approach, the deconvolved trace 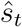 is normalized by its Fano factor. Either of these normalizations leads to similar performance in terms of encoding or decoding accuracy (data not shown).

##### Bernoulli model

The Bernoulli model can be considered as a special case of the ZIG model, by collapsing the positive responses into a delta function at 1. The responses are first binarized by thresholding; as discussed in the main text, we explore a range of different *s*_*min*_ values in the deconvolution step, and set the binarization threshold equal to *s*_*min*_. As in the ZIG model, we define 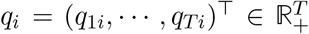 as the non-zero probability for neuron *i*.

##### Gamma model

For completeness, we also fit a gamma model to the deconvolved responses. (Note that the gamma distribution can not capture the strong bimodality that we typically observe in 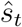.) The gamma distribution exhibits a singularity at 0 when the shape parameter *k* is less than 1. To avoid this issue, we slightly shift the observations away from 0 by adding a small positive number *ϵ* (*ϵ* = 10^−4^) to 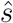 before fitting the gamma(*k, a*) model. As in the ZIG model, we denote 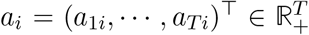 as the scale parameters for neuron *i*.

##### Parameter estimation

We estimate the parameters of the above models via maximum (pseudo-)likelihood. Details appear in Section 4.3.

#### 4.2 In vivo datasets

Two in vivo datasets are analyzed here. Both datasets are about 15 minutes long; in each case GCaMP6f was utilized as the calcium indicator. Traces are extracted using the CNMF-E software described in (Zhou *et al.*, 2018).

The first dataset is from area ADN of mouse thalamus. During stereotaxic surgery male B6/C57j mice were injected in ADN with the viral vector AAV9-hSyn-GCaMP6f (Molecular Tools Platform, Laval University). These mice were then implanted with a GRIN relay lens that was 500 microns in diameter and 4.0 mm in length (Inscopix, Inc.). The lens was positioned such that the bottom surface of the lens terminated just dorsal to the ADN. Baseplates used to attach the miniaturized fluorescent imaging endoscope (‘UCLA Miniscope’, miniscope.org) were cemented to the skull and imaging was performed using miniscopes while following the guidelines on the miniscope.org website. Recording sessions were conducted on a plus-maze (with each arm being 70cm long and 7.5 cm wide) in which animals were trained to alternate between arms. A webcam mounted above the maze tracked the position of a green and red light emitting diode that were attached to the miniscope. These were used to determine position and head direction of the mouse. Images were acquired at 30Hz. All experimental procedures followed the guidelines approved by the McGill University Animal Care Committee.

The second dataset is from area CA1 of mouse hippocampus. This dataset was collected using 2-photon imaging, while the head fixed mouse was running for a stably placed hidden (non-cued) water reward on a 2 meter belt containing discrete tactile landmarks as in (Zaremba *et al.*, 2017). Images were acquired at 60Hz (post hoc temporally decimated to 30Hz).

#### 4.3 Fitting encoding models to the data

We use a similar maximum likelihood-based fitting procedure for both the simulated data and the two real datasets. We denote *θ* = (*θ*_1_, …, *θ*_T_)^*T*^ ∈ (−180, 180]. We split the data into 60% training data, 20% validation data, and 20% test data, for two simulation datasets and ADN data. For the CA1 data, only the data from running state are used, and the data are split it into 70% training data, 10% validation data, and 20% test data.

##### ZIG model

There are three parameters, i.e., the scale parameter *a* and shape parameter *k* for the gamma component, and the probability of non-zero responses *q*. We parameterize the scale *a* and probability of non-zero responses *q* as a function of stimulus (sin(*θ*), cos(*θ*)) using neural networks, i.e.,

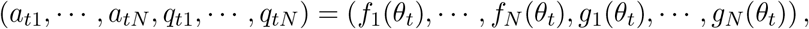

where *f* = (*f*_1_, …, *f*_N_), *g* = (*g*_1_, …, *g*_*N*_) are the output layer for *a* and *p* respectively. We use 2 hidden layers, each with tanh non-linearity, in the neural network. For the output layer, we use a logistic link function for *f* and an exponential link function for *g*. 30 nodes in hidden layers are used for the two simulated datasets and the ADN data; 15 nodes are used for the hippocampal CA1 data. We fix the shape parameter *k* to be a constant for individual neurons (i.e., *k* is neuron-dependent but not *θ*-dependent).

We optimize all the parameters by maximizing the log-likelihood using a variant of gradient descent, i.e., Adam (Kingma and Ba, 2014). Specifically, the objective function can be expressed as

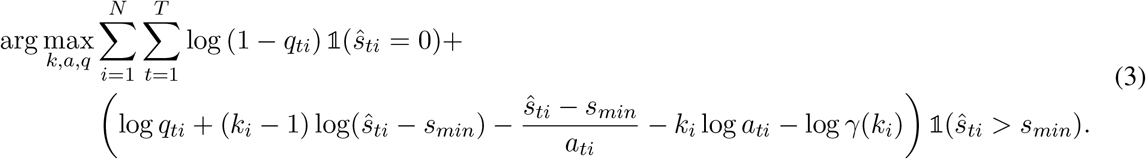

##### Poisson model

We parameterize the Poisson mean λ using a neural network with the same structure as described for the ZIG model above, with an exponential link function in the output layer. Note that for the Poisson model, the likelihood function is not a proper likelihood because the the Poisson density can not be evaluated for non-integer values. We use a “psuedo-likelihood” function instead for the maximum likelihood estimation:

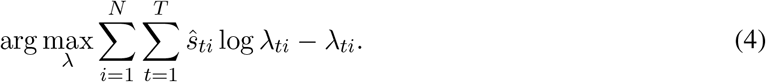

##### Bernoulli model

The probability of positive response *q* is parameterized using a neural network with the same structure as in the ZIG model, with a logistic link function in the output layer. Formally, the objective function can be defined as

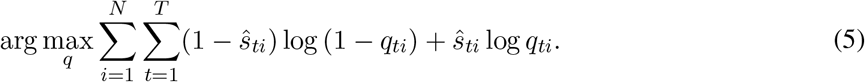

##### Gamma model

The scale parameter *a* and the shape parameter *k* are parameterized with the same neural network structure as in the ZIG model, except using an exponential link in the output layer. The objective function can be expressed as

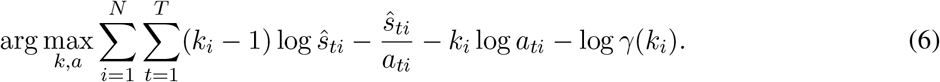

## Footnotes

1 As we will see below, it is critical to use at least a two-parameter distributional family for the slab, to capture changes in the mean and variance. The gamma family is a convenient two-parameter family that provides a good fit to the data, but other distributional families beyond the gamma could also be suitable here.

